# Centromeric DNA destabilizes H3 nucleosomes to promote CENP-A deposition during the cell cycle

**DOI:** 10.1101/215624

**Authors:** Manu Shukla, Tong Pin, Sharon A. White, Puneet P. Singh, Angus M. Reid, Catania Sandra, Alison L. Pidoux, Robin C. Allshire

## Abstract

Active centromeres are defined by the presence of nucleosomes containing CENP-A, a histone H3 variant, which alone is sufficient to direct kinetochore assembly. Once assembled at a location CENP-A chromatin and kinetochores are maintained at that location though a positive feedback loop where kinetochore proteins recruited by CENP-A itself promote deposition of new CENP-A following replication. Although CENP-A chromatin itself is a heritable entity, it is normally associated with specific sequences. Intrinsic properties of centromeric DNA may favour the assembly of CENP-A rather than H3 nucleosomes. Here we investigate histone dynamics on centromeric DNA. We show that during S-phase histone H3 is deposited as a placeholder at fission yeast centromeres and is subsequently evicted in G2 when we detect deposition of the majority of new CENP-A^Cnp1^. We also find that centromeric DNA has an innate property of driving high rates of turnover of H3 containing nucleosomes resulting in low nucleosome occupancy. When placed at an ectopic chromosomal location in the absence of any CENP-A^Cnp1^ assembly, centromeric DNA retains its ability to impose S-phase deposition and G2 eviction of H3, suggesting that features within this DNA program H3 dynamics. As RNAPII occupancy on this centromere DNA coincides with H3 eviction in G2, we propose a model in which RNAPII-coupled chromatin remodelling promotes replacement of H3 with CENP-A^Cnp1^ nucleosomes.

## Introduction

Centromeres are the defined locations on chromosomes where kinetochores are assembled and which ensure accurate chromosome segregation. In many species, centromere chromatin is distinguished by the presence of CENP-A (also known as cenH3; CID in *Drosophila*, Cse4 in *Saccharomyces*, Cnp1 in *Schizosaccharomyces*), a histone H3 variant, which substitutes for canonical H3 in specialized nucleosomes that form the foundation for kinetochore assembly [1]. The point centromeres of budding yeast (*Saccharomyces cerevisiae*) are unusual in that they are entirely DNA sequence dependent because sequence specific DNA binding proteins direct CENP-A^Cse4^ and kinetochore assembly [2]. In contrast, regional eukaryotic centromeres (human, fruit fly, plant and fission yeast) assemble CENP-A nucleosomes across extensive DNA regions that are often repetitive [3].

CENP-A is critical in defining where kinetochores are assembled since its artificial recruitment to non-centromeric chromosomal locations is sufficient to mediate kinetochore assembly [4-6]. Centromere position is normally stable, however, chromosome rearrangements that delete a normal centromere allow the appearance of neocentromeres at unusual chromosomal locations [7-9]. Moreover, dicentric chromosomes with two centromeres can be stabilised by the inactivation of one centromere without DNA loss [10,11]. Such observations indicate that CENP-A incorporation and thus centromere positioning exhibits epigenetic plasticity [12,13].

Despite the flexibility associated with CENP-A and thus centromere location, neocentromeres are rare and centromeres usually remain associated with specific DNA sequences [3,7,8,14,15]. However, despite the conservation of CENP-A and many kinetochore proteins, the underlying centromere DNA is highly divergent [16]. Nevertheless, these centromere sequences allow *de novo* CENP-A and kinetochore assembly following their introduction as naked DNA into cells [17,18]. Such analyses indicate that centromere DNA is a preferred substrate for CENP-A assembly. The CENP-B DNA binding protein somehow designates mammalian satellite repeats for CENP-A assembly. However, the mechanisms which promote assembly of CENP-A rather than H3 nucleosomes remain largely unknown [18].

During replication, parental nucleosomes are distributed to both sister-chromatids and new nucleosomes assemble in the resulting gaps by a replication-coupled process. Consequently, half of the histones in nucleosomes on G2 chromatids represent ‘old’, pre-existing subunits while the other half are newly synthesized histones incorporated during replication [19]. Measurement of CENP-A levels at vertebrate and *Drosophila* centromeres shows that they are reduced by half during replication [20,21]. Thus, CENP-A levels must be replenished each cell cycle outside of S-phase. Various analyses reveal that in contrast to canonical H3, new CENP-A is incorporated in a replication-independent process confined to a specific portion of the cell cycle. The timing of CENP-A incorporation varies between organisms, cell types and developmental stages. In mammalian cultured cells and *Drosophila* somatic tissues, new CENP-A is deposited at centromeres in late telophase/early G1 [22,23]. However, new CENP-A^CID^ is incorporated at *Drosophila* centromeres in cultured cells at metaphase and during anaphase in early embryos [21,24] whereas it is loaded during G2 in plant tissues [25].

The above studies reveal that some cells initiate chromosome segregation with a full complement of CENP-A at centromeres while others carry only half the maximum amount and replenish CENP-A levels only after mitotic entry, between metaphase and G1. Nevertheless, the key shared feature is that new CENP-A incorporation is temporally separated from bulk H3 chromatin assembly during S-phase. From S-phase until the time of new CENP-A deposition placeholder H3 nucleosomes might be temporarily assembled in place of CENP-A, or gaps completely devoid of nucleosomes may be generated at centromeres [3,26,27]. Analysis of human centromere chromatin fibers suggests that H3.3 is deposited as a placeholder in S-phase that is later replaced by new CENP-A [28]. However, the cell cycle dynamics of H3 relative to CENP-A have not been explored in substantial detail at other more tractable centromeres. Moreover, cell cycle specific replacement of H3 with CENP-A nucleosomes may be directly associated with HJURP/Mis18 mediated CENP-A deposition [29-31]. Alternatively, processes such as transcription, known to induce histone exchange [32], might aid CENP-A deposition by facilitating H3 eviction prior to or coincident with CENP-A deposition. Transcription has been observed at centromeres and is implicated in CENP-A deposition in several systems [33-42].

Once established, CENP-A chromatin has an innate ability to self-propagate through multiple cell-divisions. Such persistence is ensured by associated factors which recognize pre-existing CENP-A nucleosomes and mediate assembly of new CENP-A particles nearby [43-45]. However, the features that distinguish normal centromere DNA as being the preferred location for de novo CENP-A chromatin assembly remain unknown, although DNA binding factors such as CENP-B appear to be involved [18].

Fission yeast, *Schizosaccharomyces pombe*, centromeres are regional and have a distinct advantage in that CENP-A^Cnp1^ nucleosomes and kinetochores are assembled over specific central domains of ~10 kb that are flanked by H3K9me heterochromatin repeats [46,47]. The unique central CENP-A domain of *cen2* allows detailed analyses unencumbered by problematic repetitive centromere DNA [17,48]. Initial microscopic and genetic analyses indicated that cell cycle loading of fluorescently-tagged CENP-A at the *S.pombe* centromere cluster is either biphasic, occurring both in S-phase and G2 [49] or mid-late G2 [50]. However, the dynamics of CENP-A, H3 and RNAPII association have not been examined through the cell cycle at an individual centromere in any system.

Here we demonstrate that histone H3 is incorporated at *S.pombe* centromeres during S-phase where it serves as an interim placeholder prior to its replacement by new CENP-A during G2. This cell cycle regulated program occurs independently of CENP-A and kinetochore assembly since H3 exhibits similar cell cycle dynamics on ectopically located centromere DNA devoid of CENP-A. Moreover, ectopic centromere DNA exhibits intrinsically low H3 nucleosome occupancy and rapid nucleosome turnover. Thus H3 nucleosomes assembled on centromere DNA are intrinsically unstable. Elongating RNAPII transiently accumulates on this centromere DNA during G2, coincident with the time of H3 eviction. We propose that centromere DNA drives a program of cell cycle-coupled events ensuring the temporally regulated replenishment of CENP-A.

## Results

### New CENP-A^Cnp1^ is deposited at centromeres during G2

Previous single cell analyses indicated that CENP-A^Cnp1^ levels at the *S.pombe* centromere cluster decline during replication and are replenished during G2[50], while genetic analyses suggested that incorporation occurs during both S and G2 phases [49]. *S.pombe* CENP-A^Cnp1^ transcript levels increase prior to replication, in advance of general histone gene induction [51]. To accurately distinguish between newly synthesized and pre-existing old CENP-A^Cnp1^ protein we used Recombination Induced Tag Exchange (RITE; [52]). All pre-existing ‘old’ CENP-A^Cnp1^ was tagged with the HA epitope whereas β-estradiol induced nuclear import of Cre-EBD in *cdc25-22*/G2 arrested cells mediated recombination between Lox sites so that any ‘new’ CENP-A^Cnp1^ expressed in the following G1/S was tagged with the T7 epitope (Figure 1A and S1A-D). Following release from G2 (36→25^o^C shift) the cell population underwent synchronous cell division as indicated by a peak in septated cell frequency (cytokinesis; 60%) after ~90 minutes. S-phase in *S.pombe* coincides with cytokinesis and is followed immediately by the next G2 [53].

**Figure 1.**
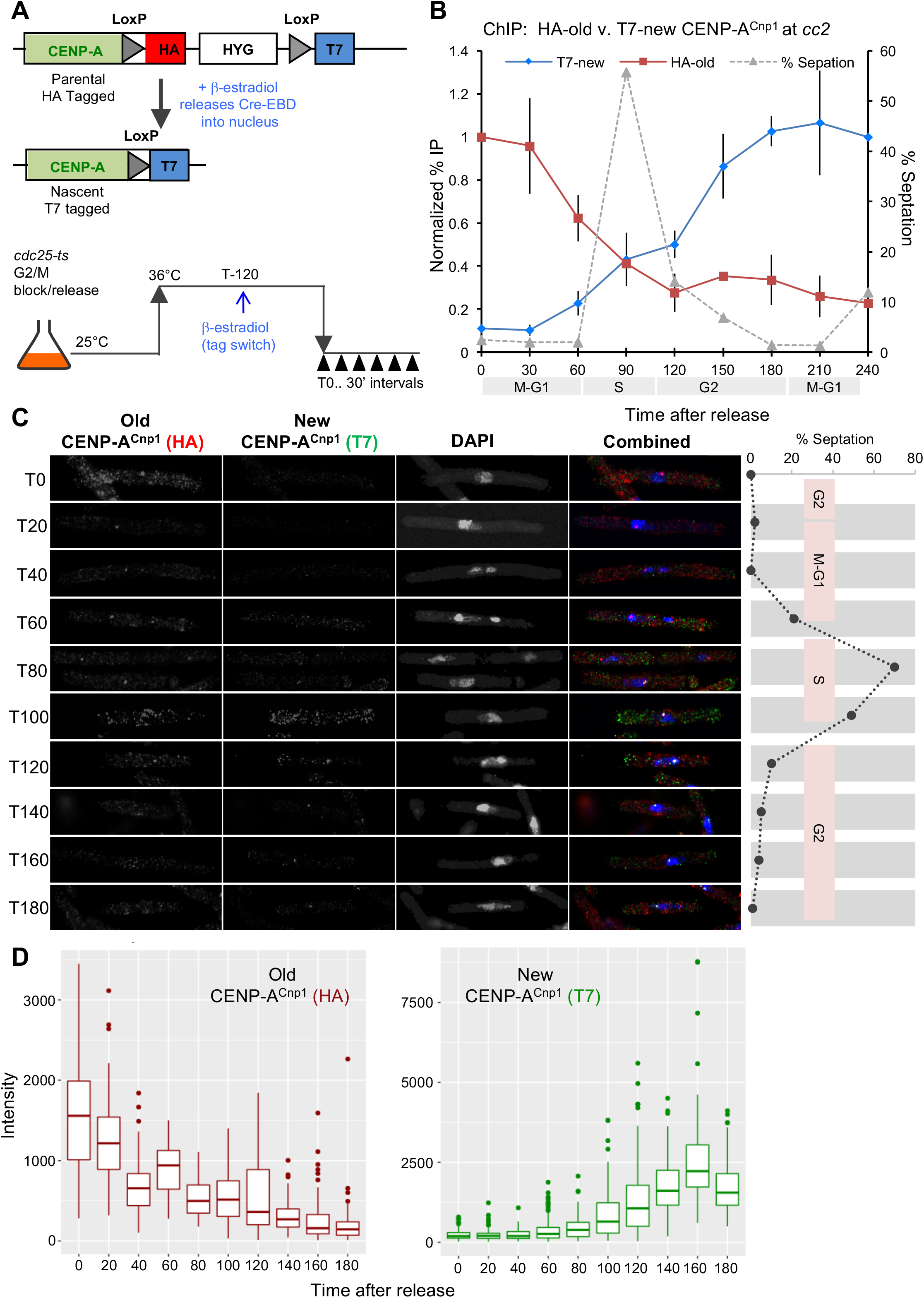
New CENP-A^Cnp1^ is deposited at centromeres during G2. (A) Diagram of <u>R</u>ecombination <u>I</u>nduced <u>T</u>ag <u>E</u>xchange (RITE) system and CENP-A^Cnp1^-RITE tag swap. *cdc25-22* ts mutant cells were blocked in G2 by incubation at 36°C, tag swap was induced by β-estradiol addition. Cells were released synchronously into the cell cycle by shifting to 25°C. Samples were collected at indicated time-points (T0-T240). (B) qChIP analysis showing the profiles for HA-tagged old and T7-tagged new CENP-A^Cnp1^ during the cell cycle at *cc2*. %IP values were normalized to values at T0 for CENP-A^Cnp1^-HA and T240 for CENP-A^Cnp1^-T7, respectively. Error bars: mean ± SD (n=3). % septation and cell cycle stages as indicated. (C) Immunolocalization to assess timing of new CENP-A^Cnp1^-T7 deposition. Representative images from each time-point are shown. Septation index and cell cycle stages as indicated. (D) Quantitation of old CENP-A^Cnp1^-HA and new CENP-A^Cnp1^-T7 intensities in individual cells (from C), n=100 for all time-points except T80 (n=94). Horizontal bars: median values, ± SD, outliers shown.

qChIP analyses at several positions across the unique central domain (*cc2*) at centromere 2 (*cen2*) revealed a drop in old CENP-A^Cnp1^-HA levels during S-phase, consistent with its dilution by distribution to both sister-centromeres. Association of new CENP-A^Cnp1^-T7 with *cc2* rose to a plateau during the subsequent G2 indicating that most new CENP-A^Cnp1^ is incorporated during G2 (Figure 1B and S1E). Microscopic analyses showed that following release from the *cdc2-22*/G2 block old CENP-A^Cnp1^-HA was detectable at centromeres throughout the time course. In contrast, new CENP-A^Cnp1^-T7 centromere localization was only detected during the next G2 (T100; Figure 1C and D).

Thus, pre-existing CENP-A^Cnp1^ declines at *S.pombe* centromeres during replication and new CENP-A^Cnp1^ is primarily incorporated in mid-late G2. Fission yeast centromeres must undergo mitosis with a full complement of CENP-A^Cnp1^ chromatin that is halved during their replication. The net loss of CENP-A^Cnp1^ from sister-centromeres may result in an increase in the size or numbers of inter-nucleosomal gaps between CENP-A^Cnp1^ nucleosomes. Alternatively, H3 containing nucleosomes may be assembled as temporary placeholders at centromeres during S-phase by replication-coupled mechanisms.

### CENP-A profiles reveal widespread deposition and distinct states during the cell cycle

We next performed ChIP-seq to assess the distribution of old-HA and new-T7 tagged CENP-A^Cnp1^ at centromeres, and genome-wide, through the cell cycle in *cdc25-22* synchronized cells. Following release from G2 and RITE-tag exchange, samples were collected every 25 minutes. At T25 old-HA CENP-A^Cnp1^ was detected across *cc2* in a series of peaks (~20) with relatively shallow intervening troughs (Figure 2A), no significant new CENP-A^Cnp1^-T7 was detected within centromeres in these M/G1 cells. As the cells proceeded through replication (peak septation/T100) both old CENP-A^Cnp1^-HA and new CENP-A^Cnp1^-T7 peaks within *cc2* appeared more distinct with deeper troughs suggesting that the positioning of CENP-A^Cnp1^ containing particles becomes more confined (Figure 2A). The majority of new CENP-A^Cnp1^-T7 was deposited in G2 and the distinctive ‘shark-tooth’ S-phase pattern became less prominent (T125), after which the pattern gradually returned to a series of peaks with intervening shallow troughs suggesting restoration of a mature CENP-A^Cnp1^ chromatin state prior to the next mitosis (T200, Figure 2B). Similar dynamics were observed at other centromeres (Figure S2A and B). These data are generally consistent with qChIP analyses which detect some new CENP-A^Cnp1^ deposition at centromeres in S-phase while most new CENP-A^Cnp1^ incorporation occurred in G2 (Figure 1).

**Figure 2.**
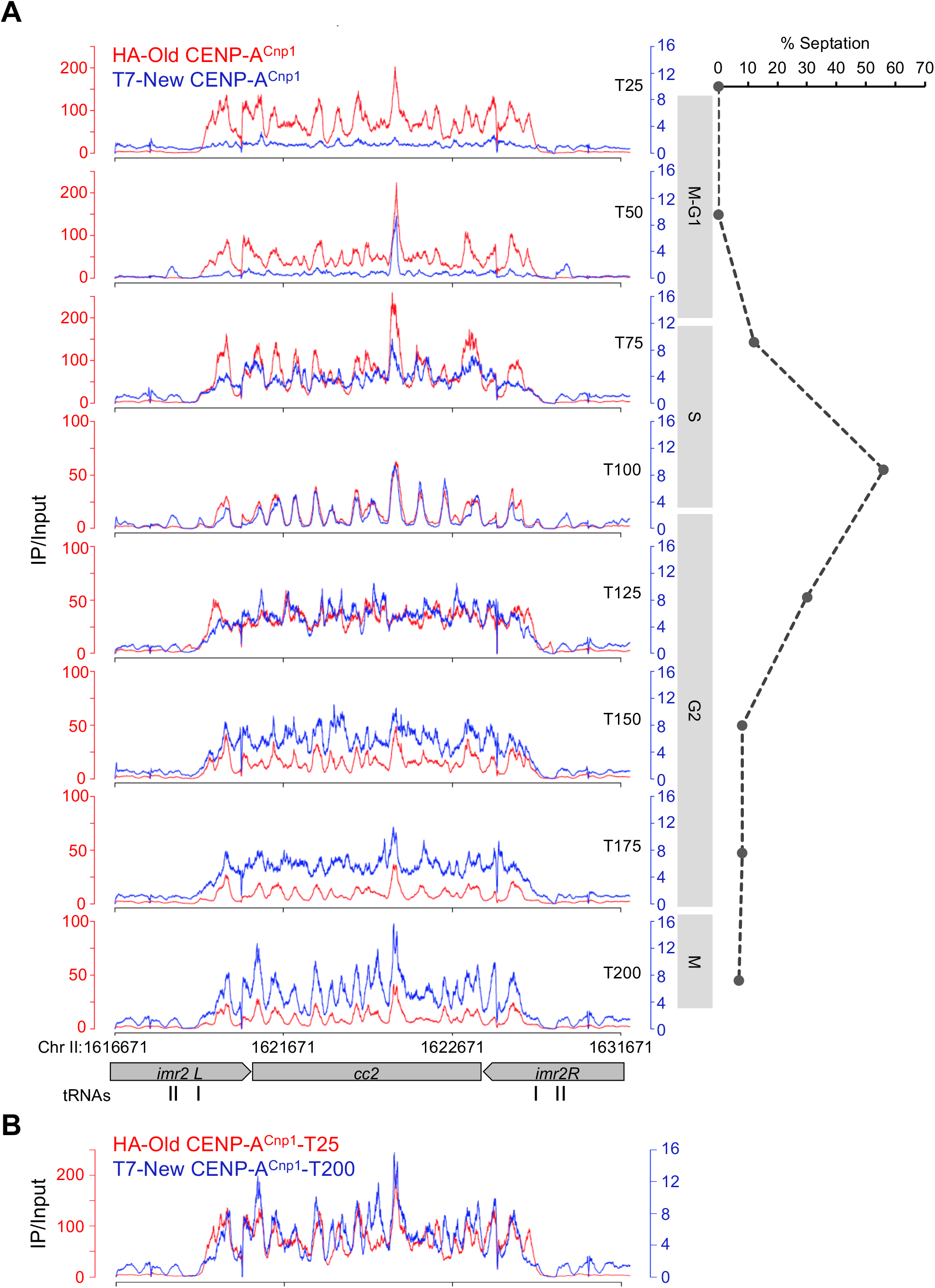
New and old CENP-A^Cnp1^ profiles reveal distinct states during the cell cycle. (A) Cell cycle ChIP-seq profiles for old CENP-A^Cnp1^-HA (red) and new CENP-A^Cnp1^-T7 (blue). Experimental scheme as in Figure 1A. Respective fold enrichment values corresponding to different time-point (IP relative to Input) for HA and T7 ChIP are shown. Cell cycle phases and chromosomal location as indicated. (B) Overlay of ChIP-Seq profiles for old CENP-A^Cnp1^-HA at T25 and new CENP-A^Cnp1^-T7 at T200.

Unexpectedly, our ChIP-Seq analysis revealed transient association of new CENP-A^Cnp1^-T7 at low levels across much of the genome prior to S-phase (T50; Figure 3A). Further analysis at this time point showed this widespread new CENP-A^Cnp1^ incorporation occurs mainly over gene bodies (Figure 3B). New CENP-A^Cnp1^ deposition within genes was most obvious when the profile was compared with that of old CENP-A^Cnp1^-HA over specific genes (Figure 3C). This widespread new CENP-A^Cnp1^ was rapidly removed as cells enter S-phase (T75), and coincided with some new CENP-A^Cnp1^-T7 accumulation within centromeres (Figure 3A). Transient incorporation of GFP-CENP-A^Cnp1^ within three genes prior to S-phase was also detected by qChIP prior to S-phase thus verifying its detection by ChIP-Seq (Figure 3D).

**Figure 3.**
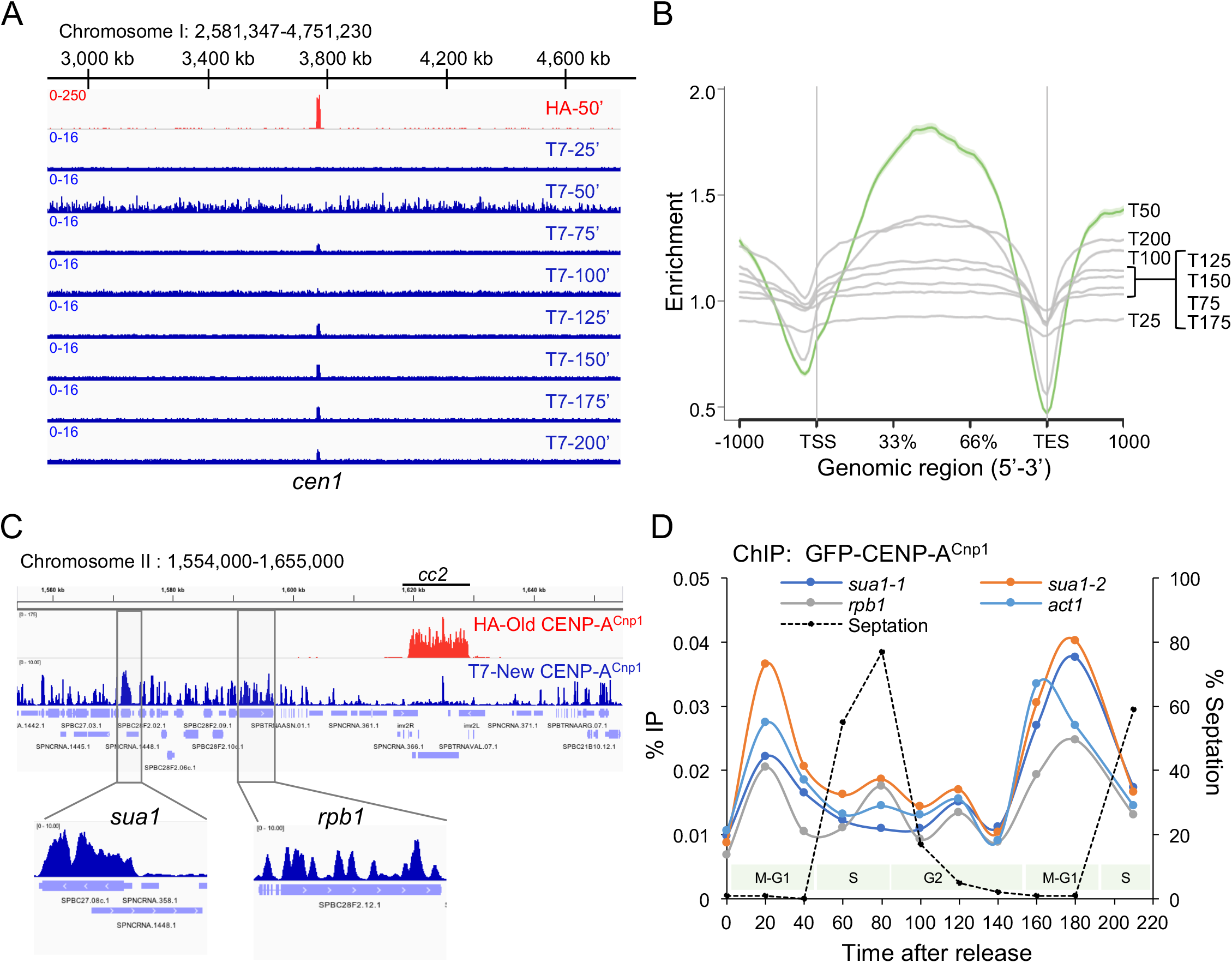
Transient association of new CENP-A^Cnp1^ throughout chromosome arms prior to S-phase. (A) Genome browser view showing cell cycle ChIP-Seq profiles of old CENP-A^Cnp1^-HA (Red: T50), new CENP-A^Cnp1^-T7 (Blue: T25-T200). Fold enrichment (IP/Input) and chromosomal location as indicated. (B) Association of new CENP-A^Cnp1^ with genes through the cell cycle. ChIP-seq enrichment values for new CENP-A^Cnp1^-T7 from indicated time-points are shown across average positions within or flanking all genes. (C) Genome browser view from chromosome II for T50 sample showing association of old CENP-A^Cnp1^-HA (Red), new CENP-A^Cnp1^-T7 (Blue). Chromosomal location of genes and *cen2* is indicated. Below: expanded profiles on exemplar *sua1^+^* and *rpb1^+^* genes showing incorporation of new CENP-A^Cnp1^-T7. (D) Representative ChIP for GFP-CENP-A^Cnp1^ cell cycle incorporation at exemplar genes (*sua1^+^, rpb1^+^ and act1^+^*). Y axis: %IP values. Septation index and cell stages as indicated.

### Histone H3 is deposited at centromeres in S-phase and evicted during G2

In many eukaryotes, CENP-A incorporation is temporally separated from replication. Chromatin fiber analyses suggest that H3.3 is deposited at human centromeres in S-phase and replaced by CENP-A in G1 [28]. A placeholder model predicts that H3 should increase within *S.pombe* centromeres during S-phase and decline when CENP-A is deposited in G2, whereas no change in H3 occupancy between S and G2 is predicted by a gap filling model (Figure 4A). To determine the interplay between H3 and CENP-A^Cnp1^ dynamics, we analyzed the levels of H3, CENP-A^Cnp1^ and H4 at *cc2* through the cell cycle in the same *cdc25-22* synchronized cell population by qChIP (Figure 4 B-D). Consistent with a placeholder model, H3 levels over *cc2* increased during S-phase and declined as cells entered G2 (Figure 4D). Reciprocally, GFP-CENP-A^Cnp1^ levels declined during S-phase but rose again to maximal levels during G2, coincident with H3 removal (Figure 4E). Importantly, the H4 levels, which report total nucleosome occupancy, remained essentially constant throughout the time course further suggesting G2 specific H3→CENP-A^Cnp1^ exchange (Figure 4F).

**Figure 4.**
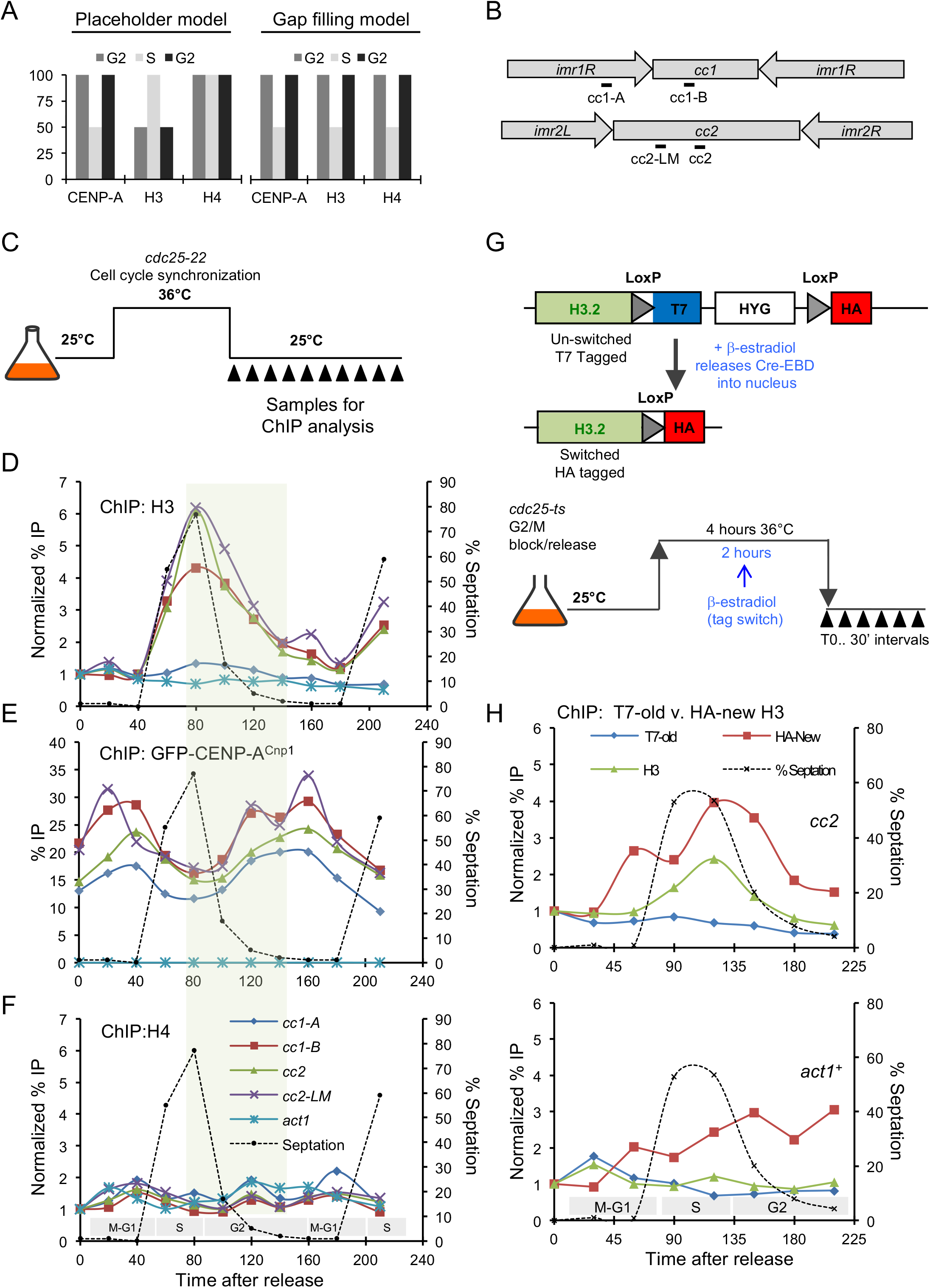
Histone H3 acts as interim placeholder for CENP-A^Cnp1^ during S-phase. (A) Illustrative graph showing expected behavior of CENP-A^Cnp1^, histone H3 and H4 at centromeres as a result of placeholder or gap filling models. (B) Schematic of *S. pombe* centromere 1 and 2 organization. Positions of primers used for qChIP as indicated. (C) Schematic of *cdc25-22* block-release cell cycle synchronization to assess cell cycle histone dynamics at centromeres. (D, E, F) Representative qChIP experiment measuring (D) histone H3, (F) H4 and (E) GFP-CENP-A^Cnp1^ levels at indicated positions at *cen1*/*cen2* (B), and on *act1^+^*, in the same samples through the cell cycle. For H3 and H4 (D and F), %IP levels were normalized first using ChIP levels at *S. octosporus act1^+^* from spiked-in chromatin and then to T0 values for each series of samples. Green rectangle: time of H3→CENP-A^Cnp1^ exchange. (G) Diagram H3.2-RITE T7-to-HA swap. The RITE cassette was integrated in-frame downstream of *hht2^+^*. Cell cycle block-release and tag swap induction as described in Figure 1A. (H) qChIP for old H3-T7, new H3-HA and total H3 levels on endogenous cc2 at *cen2* (top panel) and *act1^+^* (lower panel) through the cell cycle.

To further substantiate the timing of H3 deposition, we RITE-tagged one of the three genes (H3.2) encoding identical canonical histone H3. An assumption is that all histone H3 behaves similarly so that H3.2-RITE provides a tracer for the dynamics of all H3. In *cdc25-22*/G2 blocked cells all pre-existing old histone H3.2 was T7-tagged. Following T7-to-HA tag swap induction during the G2 block all new H3.2 was HA-tagged (Figure 4H).

Coincident with S-phase (septation peak), qChIP revealed that old H3.2-T7 levels drop within centromeres and on the constitutively expressed *act1^+^* gene by approximately half (T120, Figure 4H). This decline represents the distribution of parental H3.2-T7 nucleosomes to both sister-chromatids. New H3.2-HA accumulated within *cc2* during S/early G2 (T90-T150) but declined during mid-late G2 (Figure 4F). This decline in new H3 levels coincided with the time when most new CENP-A^Cnp1^ deposition was detected within centromeres (Figure 1 and 2). No such reduction in H3 levels was observed on the *act1^+^* gene during G2, instead new H3.2 was incorporated during replication and remained in place in the following G2. qChIP with anti-H3 on the same samples confirmed that total H3 levels increased during S-phase but declined in the following G2 within *cc2*, while little overall change occurred on *act1^+^* (Figure 4H).

These analyses suggest that H3 specifically accumulates within the CENP-A^Cnp1^-containing regions of centromeres during S-phase, but is removed during G2. New CENP-A^Cnp1^ is incorporated within these centromeric regions during G2, coincident with H3 removal. We conclude that H3 nucleosomes assembled within centromeres during S-phase serve as temporary placeholders which are replaced by CENP-A^Cnp1^ nucleosomes during G2.

### Centromere DNA alone drives histone H3 cell cycle dynamics

H3 and CENP-A^Cnp1^ deposition and eviction dynmics at centromeres may be entirely dictated by kinetochore-associated CENP-A^Cnp1^ loading factors (e.g. HJURP^Scm3^, Mis18) [31,54,55]. Alternatively, central domain sequences themselves might enforce processes that promote such dynamics. To determine if centromeric DNA programs H3 cell cycle dynamics, we utilized cells carrying 8.5 kb of *cc2* DNA at the non-centromeric *ura4* locus and with endogenous *cen2*-*cc2* replaced with *cen1* central core DNA (*cc1*) so that ectopic *cc2* (*ura4:cc2*) is the only copy of this element [35])[36](Figure 5A). qChIP on asynchronous cells confirmed that ectopic *ura4:cc2* was completely devoid of CENP-A^Cnp1^ and assembled in H3 chromatin albeit at low levels relative to *act1^+^* (Figure 5B). Remarkably, qChIP on *cdc25-22* synchronized cells revealed that H3 accumulated on ectopic *ura4:cc2* DNA during S-phase (T60-T100) but its levels subsequently fell again during G2, similar to that observed at endogenous centromeres (*cc1*; Figure 5C). We conclude that innate properties of central domain DNA promote the assembly of H3/H4 containing nucleosomes during S-phase and their removal during G2.

**Figure 5.**
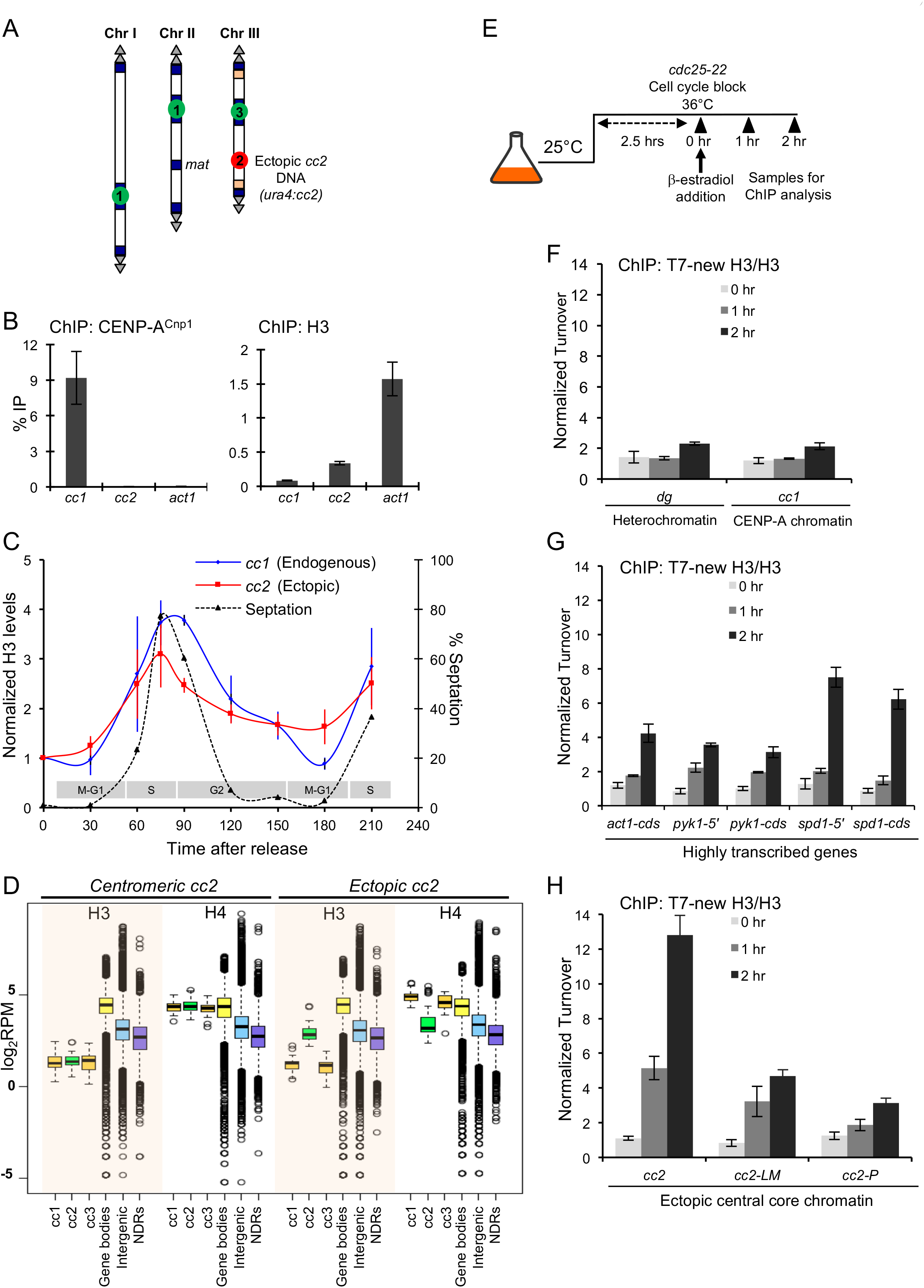
Centromere DNA destabilizes H3 nucleosome and drives histone H3 cell cycle dynamics. (A) Diagram of cells with 8.5 kb of *cc2* DNA from *cen2* inserted at an ectopic non-centromeric locus (*ura4:cc2*) and a 6 kb region of *cen2-cc2* replaced with 5.5 kb of *cen1-cc1* DNA. Allows specific ChIP on unique *ura4:cc2* which lacks CENP-A^Cnp1^. (B) qChIP for CENP-A^Cnp1^ and H3 levels at centromere located *cc1* DNA and ectopic non-centromeric *ura4:cc2* DNA. Control non-centromeric transcribed gene: *act1^+^*. Error bars: mean SD (n=3) (C) qChIP for H3 levels in *cdc25-22* synchronized cells populations at indicated time-points (T) following release into cell cycle. Septation index and cell stages as indicated. %IP values were normalized to ChIP levels at *act1^+^* and then to T0. Error bars: mean ± SD (n=3) (D) Quantitation of H3 and H4 occupancy by ChIP-Nexus. Box plot of H3 (shaded rectangles) and H4 occupancy over central core DNA (*cc1*, *cc2*, *cc3*), gene bodies, intergenic regions and NDRs in wild type cells (*centromeric cc2*) and cells carrying unique non-centromeric *ura4:cc2* (*ectopic cc2*). (E) Diagram of set-up to assess replication-independent H3 turnover. H3.2-RITE HA-to-T7 tag swap was -estradiol induced in *cdc25-22*/G2 blocked cells after two hours at 37^o^C and samples then collected 0, 1 and 2 additional hours at 37^o^C and analyzed by ChIP. (F, G, H) qChIP for new H3.2-T7 incorporated during *cdc25-22*/G2 arrest into heterochromatin *dg* repeats and *cen1 cc1* (F), highly transcribed genes (*pyk1^+^*, *spd1^+^ act1^+^*; G) and three locations with non-centromeric ectopic *ura4:cc2* (H). Normalized Turnover represents H3.2-T7 values normalized to respective total H3 levels for each sample and then to T0 value for one replicate. Error bars: mean ± SD (n=3)

### H3 nucleosomes exhibit high turnover on ectopically located centromere DNA

The cell cycle dynamics of H3 deposition and removal suggest that H3 nucleosomes assembled on ectopic central domain DNA may be inherently unstable. We next compared the steady state levels of H3 and H4 associated with native *cen2* located *cc2*, ectopic *ura4:cc2* and that detected over gene bodies (i.e. ORFs), intergenic regions (IGRs), and nucleosome depleted regions (NDRs; a signature of RNAPII promoters) using ChIP-Nexus [56]. As expected only low levels of H3 were detected over the central domain regions of endogenous centromeres (*cc1*, *cc2*, *cc3*) where most H3 is replaced by CENP-A^Cnp1^ (Figure 5D). In contrast, H4 levels across these centromeric central domains and gene bodies were equivalent. However, significantly lower levels of H3 and H4 were detected within intergenic regions, NDRs and ectopically located *ura4:cc2* DNA. These analyses suggest that H3 nucleosome assembly is strongly disfavored on ectopic *ura4:cc2* whereas CENP-A^Cnp1^ nucleosomes exhibit greater stability on *cc2* DNA within a functional centromere.

We next utilized H3.2-RITE to measure H3 turnover on ectopic *ura4:cc2* in comparison to heterochromatic repeats and highly transcribed genes in G2-arrested cells. H3.2-HA-to-T7 tag swap was induced in *cdc25-22*/G2 arrested cells and new histone H3.2-T7 incorporation monitored (Figure 5E-H). As expected, only low levels of new H3-T7 were incorporated into heterochromatin where histone turnover is low [57,58]. Consistent with transcription-coupled nucleosome exchange, high levels of new H3-T7 were incorporated into chromatin associated with the highly expressed *act1^+^*, *pyk1*^+^, *spd1*^+^ genes after one and two hours,. Intriguingly, very high levels of new H3-T7 were detected on ectopic *ura4:cc2* after just one hour indicating extensive H3 turnover. We conclude that central domain DNA must render assembled H3 nucleosomes unstable so that they are continually displaced resulting in low nucleosome occupancy.

### RNAPII accumulates on centromere DNA coincident with H3 removal

Previous analyses showed that the central CENP-A^Cnp1^ domains of *S.pombe* centromeres are transcribed and that loss of factors that promote RNAPII elongation (Clr6-CII, Ubp3 and TfIIS) enhance CENP-A^Cnp1^ deposition on naïve *cc2* DNA [35,36]. To determine if transcription, H3 turnover and CENP-A^Cnp1^ deposition on *cc2* DNA are coupled, we performed ChIP-Seq for elongating RNAPII (serine 2 phosphorylated, S2P) on *cdc25-22*/G2 synchronized cells carrying ectopic *ura4:cc2*. RNAPII-S2P association with genes whose transcription is cell cycle regulated demonstrated the expected periodic RNAPII-S2P association (Figure S3A). Thus, our synchronized cultures reliably reported cell cycle regulated RNAPII transcriptional elongation. We next examined RNAPII-S2P association with ectopic *ura4:cc2*. In G2 blocked cells (T0) relatively high levels of elongating RNAPII were detected over *ura4:cc2*, however upon release from the block, the levels of associated RNAPII-S2P declined to a minimum (T40) and accumulated again during G2 (T80-T120) (Figure 6B). RNAPII-S2P qChIP at several locations within *ura4:cc2* confirmed that elongating RNAPII occupancy falls during S-phase and increases during G2 (T100-T140; Figure 6C). The pattern of RNAPII-S2P association with *cc1* and *cc2* at endogenous centromeres was confounded by the appearance of a more dominant RNAPII-S2P association peak during mitosis (Figure 6C lower panel and Figure S3B). This M phase RNAPII-S2P peak must be kinetochore imposed as it was not detected on ectopic *ura4:cc2* (see Discussion). Nevertheless, levels of elongating RNAPII also appeared to increase at endogenous centromeres during early-mid G2 (Figure 6C and S3B) and RNAPII-S2P can be detected in affinity selected CENP-A^Cnp1^ chromatin (Figure 6D). We suggest that the more obvious elongating RNAPII accumulation on ectopic *ura4:cc2* during G2, when assembled in H3 nucleosomes only, represents an extreme version of the events that occur on centromere located *cc2* which enters G2 assembled in chromatin composed of half H3 and half CENP-A^Cnp1^ nucleosomes. We conclude that central domain DNA from *S.pombe* centromeres programs transcription-coupled H3 nucleosome destabilization during G2 resulting in their replacement with CENP-A^Cnp1^ containing nucleosomes when a sufficient route of CENP-A^Cnp1^ supply is available.

**Figure 6.**
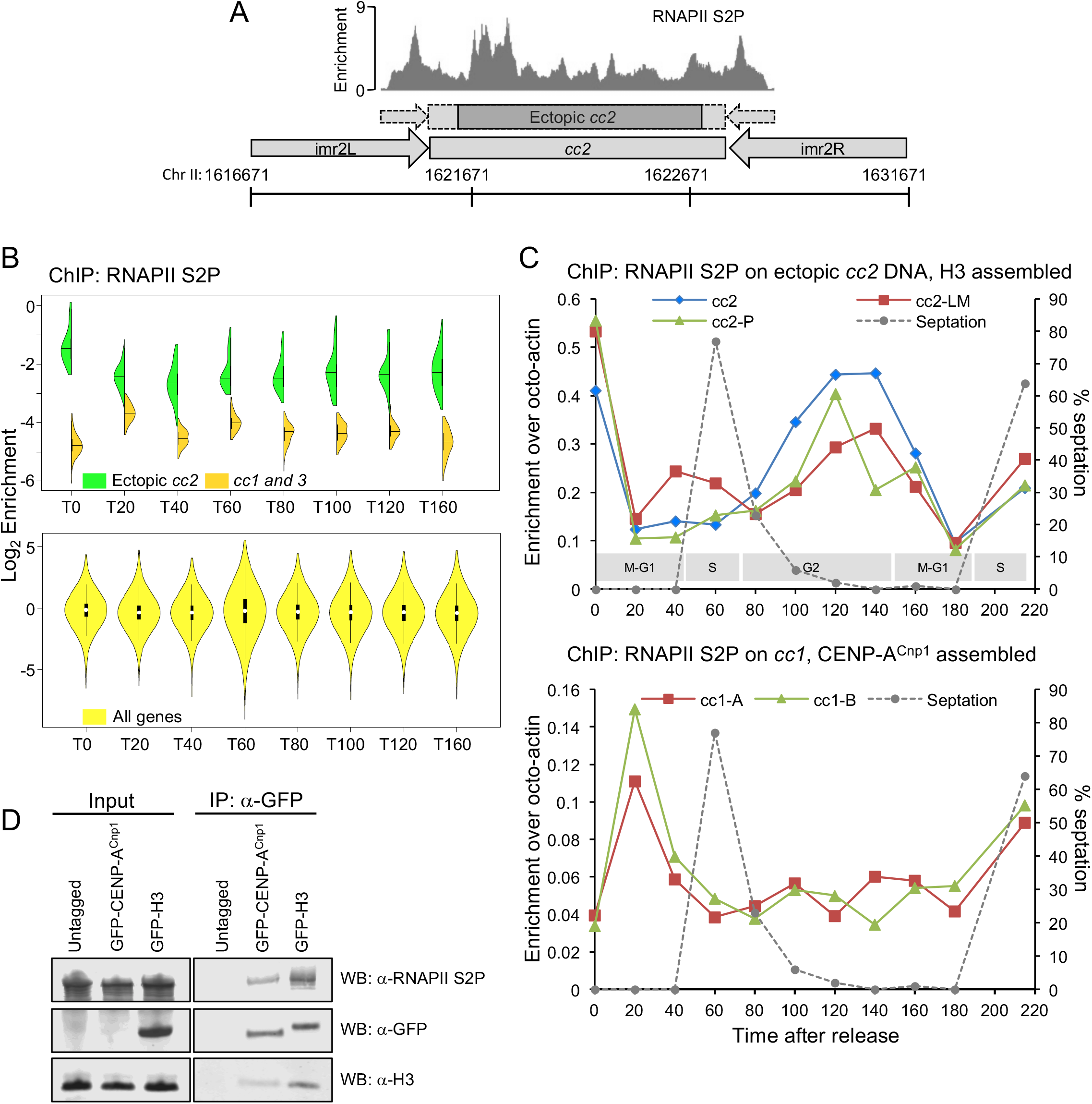
RNAPII accumulates on centromere DNA coincident with H3 removal. (A) Elongating RNAPII-S2P ChIP-Seq in *cdc25-22* synchronized cell cultures. (B) Top: RNAPII-S2P ChIP-Seq profile over ectopic *ura4:cc2* DNA is shown for a representative G2/T0 sample. Y axis shows enrichment (IP/Input). Chromosome coordinates, diagram of ectopic *cc2* and unique region within ectopic *cc2* are indicated (dark shading) (B) Violin plots of RNAPII-S2P levels over non-centromeric *ura4:cc2* (green) centromeric cc1/cc3 (orange) and genes (yellow) in T0-T160 from *cdc25-22* synchronized cell cultures. (C) qChIP for RNAPII-S2P levels on non-centromeric *ura4:cc2* (top), and centromeric *cc1* (bottom) in *cdc25-22* synchronized cell cultures (T0-T220). %IP levels were normalized using ChIP levels at *S. octosporus act1^+^* from spiked-in chromatin. Septation index and cell cycle phases as indicated. (D) RNAPII-S2P association with native affinity selected CENP-A^Cnp1^ chromatin. GFP-CENP-A^Cnp1^ or GFP-H3 chromatin was affinity selected from MNase released chromatin using anti-GFP antibody. Untagged chromatin served as a control. Anti-GFP, anti-H3 and anti-RNAPII S2P western analyses of inputs and immunoprecipitates as indicated.

## Discussion

To understand more fully how CENP-A^Cnp1^ chromatin domains are established and propagated on particular sequences across multiple generations, we focused on the cell cycle dynamics of *S.pombe* centromere-associated chromatin. RITE-tag swap experiments allowed us to determine that new CENP-A^Cnp1^ is incorporated at *S.pombe* centromeres in mid-to-late G2. Our analyses also conclusively demonstrate that histone H3 is deposited as a temporary placeholder during S-phase. Importantly, these measurements pinpoint a specific window in G2 where H3→CENP-A^Cnp1^ nucleosome exchange occurs. In addition, we show that the nascent CENP-A^Cnp1^ chromatin profile is highly dynamic exhibiting stage specific patterns through the cell cycle. Strikingly, we find that ectopically located centromere DNA assembles inherently unstable H3 nucleosomes which exhibit high turnover rates, and during replication this DNA also directs elevated incorporation of H3 nucleosomes that are evicted in the following G2 when additional elongating RNAPII is recruited. Together our analyses support a model where centromere specific DNA drives a sequence-directed program to promote H3 nucleosome eviction and the incorporation of new CENP-A^Cnp1^ nucleosomes (Figure 7).

**Figure 7.**
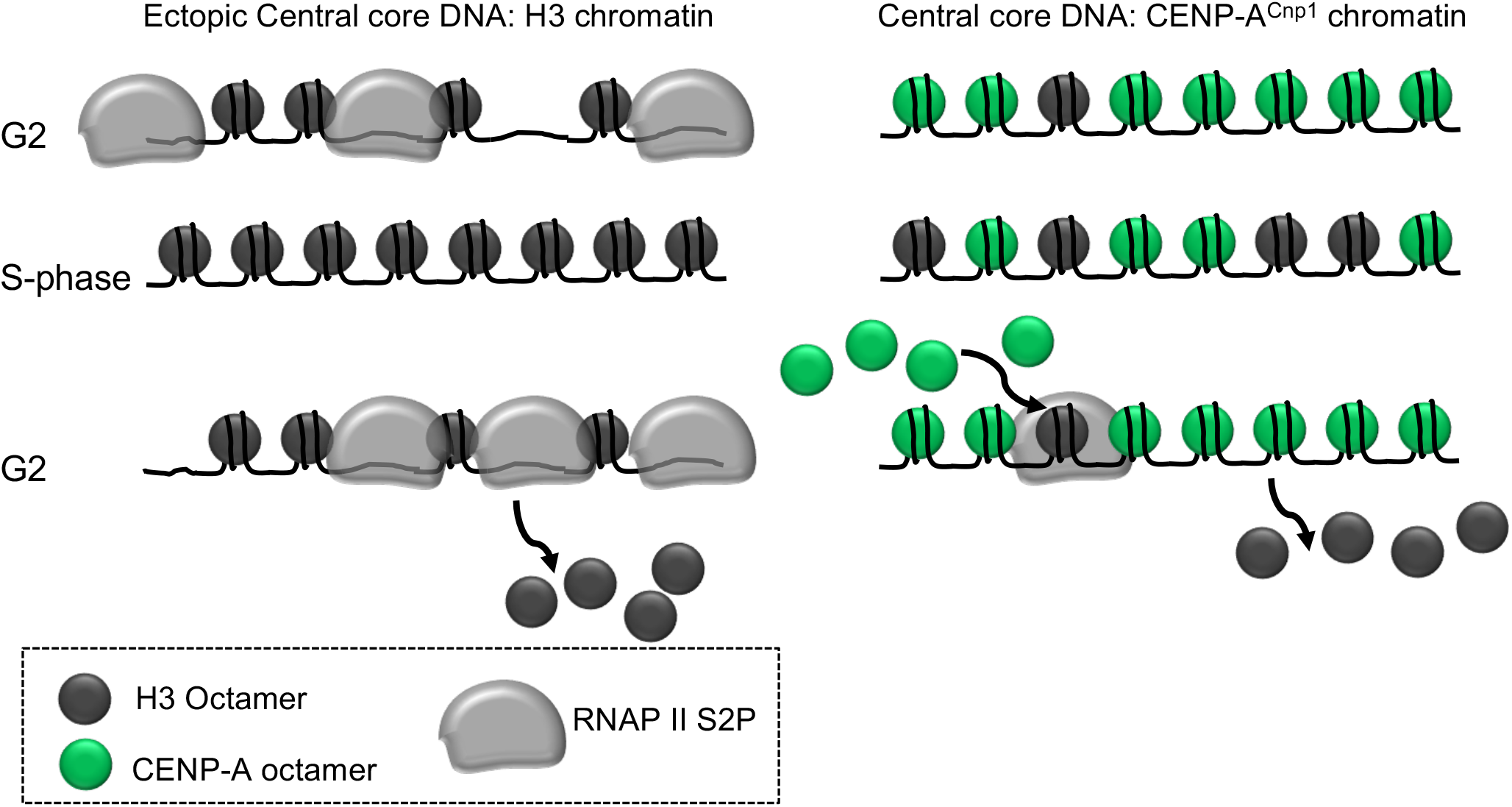
Model for centromere DNA driven histone dynamics. Left: Ectopic cc2 centromere DNA drives H3 deposition in S-phase, RNAPII recruitment and H3 eviction in G2 despite the absence of CENP-A^Cnp1^ chromatin or CENP-A^Cnp1^ dedicated deposition machinery. Right: New CENP-A^Cnp1^ is incorporated at centromeres in G2. H3 nucleosomes are transiently assembled as placeholders at centromeres during S-phase and replaced in the following G2 by new CENP-A^Cnp1^ nucleosomes. RNAPII recruited during G2 facilitates H3 nucleosomes disassembly.

The cell cycle timing of new CENP-A loading at centromeres varies between organisms; however, a conserved feature is that new CENP-A incorporation is uncoupled from replication when most new H3 chromatin assembly occurs [21-25]. Key components required for CENP-A chromatin maintenance at human centromeres (Mis18 complex, Mis18BP1^KNL2^, HJURP) are recruited to the centromeres in a temporally restricted manner prior to, or coincident with, new CENP-A deposition [5,59,60]. However, Mis18 and Scm3^HJURP^ remain associated with *S.pombe* centromeres through most of the cell cycle apart from a brief period in mitosis [31,54]. Thus, *S.pombe* centromeres might be more generally competent for new CENP-A^Cnp1^ deposition throughout the cell cycle. It was therefore critical to specifically distinguish old parental CENP-A^Cnp1^ from newly synthesized CENP-A^Cnp1^ at individual centromeres. Previous studies have suggested either biphasic (both S and G2) or G2 specific replenishment of *S.pombe* CENP-A^Cnp1^ and relied on microscopic measurement at the centromere cluster rather than association with chromatin [49,50]. RITE-tagging allowed detailed examination of the behavior of both old and new CENP-A^Cnp1^ at specific centromeres. Our data show that in *S.pombe* the majority of the new CENP-A^Cnp1^ deposition occurs after cytokinesis, which represents G2, with only low new CENP-A^Cnp1^ deposition during replication. These analyses refine previous studies of CENP-A^Cnp1^ replenishment and suggest that other mechanisms, apart from the temporally regulated recruitment of CENP-A loading factors to centromeres, can influence the cell cycle stage specific restriction of new CENP-A^Cnp1^ incorporation.

An unexpected finding was that new CENP-A^Cnp1^ is transiently incorporated within genic regions along chromosome arms prior to replication. In *S.pombe* the gene encoding CENP-A^Cnp1^ (*cnp1^+^*) is known to be expressed prior to the peak of canonical histone gene expression in advance of S-phase. As a consequence of this earlier CENP-A^Cnp1^ expression, there is a brief window during the cell cycle where the ratio of soluble CENP-A^Cnp1^ versus canonical histone H3 may be skewed. This could account for the transient widespread incorporation of CENP-A^Cnp1^ within genes which may occur through transcription-coupled nucleosome turnover events. In essence, this portion of the cell cycle exhibits similarity to cells in which CENP-A^Cnp1^ is overexpressed. Non-centromeric CENP-A incorporation occurs at many locations distributed throughout the genome [61] and when it is overexpressed [35,62,63]. Conversely, histone H3 can replace CENP-A^Cnp1^ within *S.pombe* centromeres when the H3:CENP-A^Cnp1^ ratio is perturbed [64]. The chromosomally located CENP-A^Cnp1^ that we detect is rapidly removed, conceivably through replacement with canonical H3 during replication, additional transcription-coupled turnover events or degradation. This genome ‘sampling’ by widespread CENP-A^Cnp1^ incorporation every cell cycle could potentially contribute to the formation of neocentromeres when conditions demand.

G2 specific loading of new CENP-A^Cnp1^ prompted us to also evaluate H3 and H4 cell cycle dynamics at centromeres. Comparison of the relative levels of CENP-A^Cnp1^, H3 and H4 within centromeric central domain showed that histone H3 is transiently deposited as a placeholder during S-phase and later exchanged for new CENP-A^Cnp1^ in G2. H3.3 may act as an interim placeholder for CENP-A at human centromeres [28], thus H3 placeholder function may be conserved across metazoa where new CENP-A deposition is separated from replication. The transient use of H3 as a placeholder in *S.pombe* is important as it identifies a cell cycle period where specific H3→CENP-A replacement events must take place and involve factors that mediate histone exchange or complete nucleosome turnover.

ChIP-seq analyses of old and new CENP-A^Cnp1^ at S.pombe centromere revealed unexpected patterns of CENP-A^Cnp1^ association during the cell cycle. Large scale changes in CENP-A^Cnp1^ nucleosome profiles were observed that correspond to S-phase, early-mid G2 and mitosis. It seems likely that these profiles represent various stages of CENP-A^Cnp1^ chromatin assembly perhaps corresponding to its seeding, loading and maturation. CENP-A chromatin assembly has been proposed to occur in two steps, the first involving weak association of new CENP-A with centromeric DNA followed by its actual assembly into fully formed nucleosomes [65]. Our detection of distinct CENP-A^Cnp1^ ChIP-Seq profiles during G2 and late G2/M suggests that intermediate states exist prior to the assembly of mature CENP-A^Cnp1^ nucleosomes. These different cell cycle states predict the involvement of distinct chromatin remodeling activities. Various ATP-dependent chromatin remodelers such as Hrp1^Chd1^, Fft3^Fun30^, Swr1 and Ino80 have been shown to localize within CENP-A^Cnp1^ chromatin in unsynchronized fission yeast cell populations [66-68] (Kulasegaran-Shylini et al., submitted). Further detailed analyses are required to determine which remodeling factors are responsible for the transitions between these distinct chromatin states and their role in preserving centromere integrity. Similar cell cycle profiling of nucleosomes at non-repetitive endogenous or neocentromeres in other organisms will determine if such features are generally conserved.

Where tested, centromeric DNA sequences are clearly a preferred substrate for *de novo* CENP-A assembly [48, 69, 70]. The embedded features that identify centromere DNA for efficient CENP-A assembly involves DNA binding factors such as CENP-B and processes such as transcription [33-42]. Transcription is a potent chromatin remodeling mechanism and is coupled to histone exchange; new histone H3.3 is deposited within transcribed genes and H2A is exchanged for the variant H2A.Z in the highly dynamic NDR promoter-proximal nucleosomes of many organisms [71]. Our finding that elongating RNAPII-S2P increases on S*.pombe* centromere DNA at the time that correlates with H3 eviction and CENP-A^Cnp1^ incorporation suggests that transcription-coupled remodeling events may define this centromeric DNA by driving H3→CENP-A exchange. Consistent with this is the presence of a high density of transcriptional start sites within this centromere DNA that may promote pervasive low quality transcription [36].

The association of elongating RNAPII was less evident during G2 at endogenous centromeres than on ectopically located central domain DNA (*ura4:cc2*). Ectopic central domain DNA lacks CENP-A^Cnp1^ nucleosomes whereas at centromeres it is assembled in half H3 and half CENP-A nucleosomes when cells enter G2. The CENP-A N-terminal tail is distinct and lacks key lysine residues (K4, K36) whose modification in H3 aids transcription. It is therefore not surprising that lower elongating RNAPII levels are observed at endogenous centromeres than on ectopic centromere DNA. Indeed, CENP-A chromatin has been shown to inhibit transcription *in vitro* [72]. It is likely that limited transcription-coupled turnover also contributes to H3→CENP-A exchange at endogenous centromeres.

Our analyses demonstrate that fission yeast centromeric DNA has an intrinsic ability to recruit elongating RNAPII and destabilize H3 nucleosomes in a cell-cycle regulated manner. Such embedded features must earmark these sequences for the efficient *de novo* establishment of CENP-A^Cnp1^ chromatin and kinetochores. Once established, CENP-A/kinetochore associated assembly factors (Mis18, HJURP) ensure CENP-A^Cnp1^ maintenance at these locations. We identify events that precede and accompany the replenishment of new CENP-A^Cnp1^ at fission yeast centromeres during the cell cycle. It is now essential to determine whether centromeric DNA in other organisms also have an innate capacity to drive H3 eviction in favor of CENP-A.

## Acknowledgements

We are grateful to Fred van Leeuwen for RITE system constructs. We would like to thank Nick Toda, Pauline Audergon, Ryan Ard, Daniel Robertson and David Kelly for their useful suggestions and technical inputs. M.S. was partly supported by an EMBO Long Term Fellowships (ALTF 664-2011). P.T was partly supported by funding from the European Commission Network of Excellence EpiGeneSys (HEALTH-F4-2010-257082) to RCA. S.C was supported by a Wellcome PhD Studentship [086574]. R.C.A. is a Wellcome Principal Research Fellow (200885); the Wellcome Centre for Cell Biology is supported by core funding from Wellcome (203149).

## Author Contributions

M.S and R.C.A jointly conceived the study. M.S, P.T, S.A.W., P.P.S, A.M.R, and S.C performed experiments. M.S, P.T and A.L.P analysed data. M.S and R.C.A wrote the manuscript with inputs from other authors.

## Materials and Methods

### Cell growth and manipulation

Standard genetic and molecular techniques were followed. Fission yeast methods were as described [73]. YES (Yeast Extract with Supplements) was used as a rich medium or PMG (Pombe Minimal Glutamate) for growth of cells in liquid cultures. 4X YES was used for experiments where higher cell numbers were required. CENP-A^Cnp1^ and H3.2 RITE strains were constructed by PCR amplifying HA/T7 or T7/HA RITE cassettes described in [52] and integration at the endogenous gene locus. Cre-EBD open reading frame was PCR amplified from pTW040 [52] and cloned in pRAD11, 13, 15 vectors containing different strengths of ADH promoters (Gift from Y. Watanabe). The Cre-EBD plasmids were integrated at the *ars1* locus by transformation of the plasmids DNAs linearized by *Mlu*I digestion. Strains are described in Table S1.

### Cytology

Cells were fixed with 3.7% formaldehyde for 7 minutes at room temperature. Immuno-localization staining was performed as described [74]. The following antibodies were used at 1:100 dilution: Anti HA (Abcam, ab9110), anti-T7 (Merck, 69522); Alexa 594 and 488 labelled secondary antibodies at 1:1000 dilution (Life Technologies). Single images were acquired with a Zeiss LSM 880 confocal microscope equipped with Airyscan superresolution imaging module, using a 100X/1.40 NA Plan-Apochromat Oil DIC M27 objective lens. Images were processed using ZEN Black image acquisition and processing software (Zeiss MicroImaging). Images were analysed using Image J.

### ChIP-qPCR

Cells were fixed with 1% formaldedyde for 15 minutes followed by quenching with 250 mM Glycine for 5 minutes at room temperature. ChIP was essentially performed as described [64] using antibodies against CENP-A^Cnp1^ (Sheep in-house; 10 μl), H3 (ab1791, Abcam; μ2l), H4 (Merck, 05-858; μ2l), HA (12CA5, in-house preparation; μ2l), T7 (Abcam, ab9138; μ2l), GFP (ThermoFisher Scientific, A11122; μ2l) and Phopho serine 2 RNA polymerase II (Abcam, ab5095; μ2l). 2.5×10^8^ cells were used per ChIP sample. Quantitative PCR reactions were performed in 10 μl volume with Light Cycler 480 SybrGreen Master Mix (Roche). The data were analysed using Light Cycler 480 Software 1.5 (Roche). q-PCR primers are listed in Table S2.

### ChIP-Seq and ChIP-Nexus

Due to higher number of cells required, for ChIP-Seq from synchronized cell cultures, cells were grown in 4X-YES and ChIP protocol was modified. Briefly, cell pellets corresponding to 7.5×10^8^ cells were lysed by four 1 minute cycles of bead beating in 500 μl of lysis buffer (50 mM HEPES-KOH, pH 7.5, 140 mM NaCl, 1 mM EDTA, 1% Triton X-100, 0.1% sodium deoxycholate). Insoluble chromatin fraction was isolated by centrifugation at 6000*g*. The pellet was washed with 1 ml lysis buffer. This washed pellet was gently resuspended in 300 μl lysis buffer containing 0.2% SDS and sheared by sonication with Bioruptor (Diagenode) for 30 minutes (30 s On, 30 s off at high setting). 900 μl of lysis buffer (without SDS) was added and samples were clarified by centrifugation at 17000g for 20 minutes. Supernatants were used for ChIP. Respective antibody and protein G-dynabeads (ThermoFisher Scientific) amounts were scaled up according to the cell number. Immunoprecipitated DNA was recovered using Qiagen PCR purification kit. ChIP-Seq libraries were prepared with 1-5 ng of ChIP or 10 ng of input DNA. DNA was end-repaired using NEB Quick blunting kit (E1201L). The blunt, phosphorylated ends were treated with Klenow exo^−^ (NEB, M0212S) and dATP to yield a protruding 3-’A’ base for ligation of NEXTflex adapters (Bioo Scientific) which have a single ’T’ base overhang at the 3’ end. After adapter ligation DNA was PCR amplified with Illumina primers for 13-15 cycles and library fragments of ~300 bp (insert plus adaptor sequences) were selected using Ampure XP beads.

ChIP-Nexus libraries were prepared essentially as described [56]. Briefly, protein G-dynabeads bound DNA-protein-complexes were affinity selected using antibodies. DNA was end repaired using T4 DNA polymerase (NEB, M0203S), DNA polymerase I large fragment (NEB, M0210S) and T4 polynucleotide kinase (NEB, M0201S). A single 3’-A overhang was added using Klenow exo^−^ polymerase. Adapters were ligated and blunted again by Klenow exo^−^ polymerase to fill in the 5’ overhang first and then by T4 DNA polymerase to trim possible 3’ overhangs. Blunted DNA was then sequentially digested by lambda exonuclease (NEB, M0262S) and RecJf (NEB, M0264L). Digested single strand DNA was then eluted, reverse cross-linked and phenol-chloroform extracted. Fragments were then self-circularized by Circligase (Epicentre, CL4111K). An oligonucleotide was hybridized to circularized single DNA for subsequent *Bam*HI digestion in order to linearize the DNA. This linearized single strand DNA was then PCR-amplified using adapter sequences and libraries were purified and size selected using Ampure XP beads. The libraries were sequenced following Illumina HiSeq2500 work flow.

Next generation sequencing libraries were aligned to *S. pombe* build ASM294v2.20 using Bowtie2. ChIP-Seq reads with mapping qualities lower than 30, and read pairs mapped over 500-nt apart or less than 100-nt, were discarded. All the ChIP-seq data were normalized with respect to their input data. ChIP peaks were identified from the alignments using macs2 with the corresponding input data. Deeptools was used to generate genome wide enrichment profiles using a 50 bp window size and the data visualized using the IGV genome browser. ChIP-Nexus data were analysed using MACE [75].

### Co-immunoprecipitation

Cell were grown in 4X-YES and 5×10^9^ cells were used per IP. Briefly, cells were harvested by centrifugation at 3500***g***, washed twice with water and flash frozen in liquid nitrogen. Frozen cell pellets were ground using Retsch MM400 mill. The grindate was resuspended in lyisis buffer (10 mM Tris pH7.4, 5 mM CaCl_2_, 5 mM MgCl_2_, 50 mM NaCl, 0.1% IGEPAL-CA630 and supplemented with Halt protease and phosphatase inhibitor (ThermoFisher Scientific, 1861281) and 2 mM PMSF. Chromatin was solubilized by incubation with 2 units of Micrococcal nuclease (Sigma, N3755) for 10 minutes at 37° C. MNase digestion was stopped by adding EGTA to 20 mM and lysate were rotated at 4° C for 1 hour to ensure chromatin solubilization. Lysates were clarified by centrifugation at 20,000***g*** for 10 minutes and supernatants were used for immunoprecipitation. Cleared lysates were incubated with 10 μg of anti-GFP antibody (Roche, 11814460001) and 25 μl of protein G-dynabeads (ThermoFisher Scientific), which were already crosslinked with DMP (dimethyl pimelimidate) (ThermoFisher Scientific, 21666), for 1 hour at 4° C with gentle rotation. Bead-bound affinity-selected chromatin was washed three times with lysis buffer and eluted with LDS loading buffer (ThermoFisher Scientific, 84788). Western blotting detection was performed using anti-GFP (Roche, 11814460001), anti-H3 (Abcam, ab1791) and anti-phospho-Serine2-RNA polymerase II (Abcam, ab5095) and secondary IRDye® 680RD anti-Rabbit (Licor, 926-68073) and IRDye® 800CW anti-Mouse antibodies (Licor, 926-32210).

## Data accessibility

Sequencing data have been submitted to the Gene Expression Omnibus under accession no. GSE106494.

